# Label-Free Monitoring of Cancer-associated Fibroblast Activation using NADH Fluorescence Lifetime Imaging

**DOI:** 10.64898/2026.01.23.701296

**Authors:** Nina Anseeuw, Carmen Escalona-Noguero, Giel Vankevelaer, Olivier De Wever, Ilaria Elia, Susana Rocha, Guillermo Solís-Fernández

**Affiliations:** Molecular Imaging and Photonics, Department of Chemistry, KU Leuven, Belgium; Laboratory of Metabolic Regulation of Cell Function, Department of Cellular and Molecular Medicine, KU Leuven, Belgium; Laboratory of Experimental Cancer Research, Department of Human Structure and Repair, Ghent University, Ghent, Belgium

## Abstract

Cancer-associated fibroblasts (CAFs) are key regulators of tumor progression, yet their activation state is commonly assessed using static, endpoint assays that do not allow dynamic analysis in living cells. Although CAF activation is accompanied by pronounced metabolic remodeling, label-free approaches that exploit these changes for real-time monitoring remain limited. Here, we demonstrate that NADH fluorescence lifetime imaging microscopy (FLIM) provides a non-invasive readout of this process. CAFs activated with transforming growth factor beta (TGF-β) exhibit a reproducible shift toward longer NADH fluorescence lifetimes compared to non-activated cells, consistent with changes in the relative contributions of free and protein-bound NADH. By combining live-cell FLIM with α-smooth muscle actin staining in the same cells, we directly link metabolic signatures to cellular activation state. We further demonstrate the potential of this approach to dynamically monitor CAF activation in live, migrating cells. Together, these results establish NADH fluorescence lifetime imaging as a label-free metabolic approach for monitoring CAF activation dynamics, complementing conventional marker-based methods and enabling continuous monitoring of tumor-stroma interactions.

## Introduction

Cancer progression is recognized as a multicellular process shaped not only by mutations within tumor cells, but also by their interactions with the surrounding tumor microenvironment (TME). This microenvironment is composed of various non-malignant stromal cells, including endothelial cells, immune cells, and cancer-associated fibroblasts (CAFs)^1–3^. CAFs play an essential role in tumor development, metastasis and therapy resistance^3,4^, and often represent the dominant stromal cell population, actively influencing the biochemical and mechanical properties of the TME^2,3^. Unlike normal fibroblasts, which support tissue homeostasis and wound healing, CAFs are reprogrammed into an activated, tumor-promoting state^3–5^. Depending on cues from their microenvironment, CAFs can adopt different functional states, including myofibroblast-like (myCAF) or inflammatory (iCAF) phenotypes, and dynamically switch between these states^6^. One of the known drivers of the activation into myCAF phenotype is the cytokine transforming growth factor beta (TGF-β)^7,8^, which signals through both SMAD-dependent and -independent signaling pathways^7,8^. This activation leads to increased expression of classic markers of CAF activation, including α-smooth muscle actin (α-SMA), fibroblast activation protein (FAP) and plasminogen activator inhibitor-1 (PAI-1)^9,10^.

Understanding CAF activation is essential to dissect interactions between tumor cells and the TME, as well as the mechanisms that drive tumor growth and progression. CAF activation is traditionally assessed using immunohistochemical staining or Western blotting for the classical markers (α-SMA, FAP or PAI-1)^9,10^. However, these approaches are inherently endpoint measurements, as they require fixation or cell lysis, and therefore do not permit time-resolved monitoring of CAF activation dynamics in living cells. While genetically encoded reporter systems for α-SMA expression have been developed to assess fibroblast activation in living cells^11^, their use requires prior modification of the cells, which may alter cellular behavior and limit broader applicability^11^. Consequently, there remains a need for label-free, dynamic techniques that allow continuous monitoring of CAF activation over time.

Here, we propose a novel strategy to monitor CAF activation in live cells using NAD^+^/NADH autofluorescence as a readout of cellular metabolic state. In addition to transcriptional reprogramming, TGF-β mediated CAF activation involves substantial metabolic changes, including increased glycolysis, oxidative stress, and catabolic activity^12–14^. Guido et al. demonstrated that activation of the TGF-β pathway is sufficient to downregulate caveolin-1 (Cav-1) expression in stromal cells, thereby promoting processes such as autophagy and aerobic glycolysis^12^. This metabolic shift enhances the production of energy-rich metabolites like L-lactate and pyruvate, resulting in a Warburg-like phenotype characteristic of activated stromal cells^12^.

Intrinsically linked to this metabolic reprogramming are alterations in NAD^+^/NADH dynamics. NAD^+^ is reduced to NADH during glycolysis, pyruvate conversion and the TCA cycle, whereas NADH is oxidized back to NAD^+^ via oxidative phosphorylation or, under glycolytic conditions, by lactate dehydrogenase^15– 19^. In addition to these redox reactions, intracellular NAD^+^ levels are maintained through biosynthetic routes such as the NAD^+^ salvage pathway^16^. Moreover, glycolysis and the TCA cycle represent only two of many metabolic pathways that utilize NAD^+^/NADH as a redox cofactor, collectively contributing to the intracellular redox ratio. The balance between these opposing fluxes continuously shapes the intracellular redox ratio, rendering NADH a sensitive indicator of cellular metabolic state.

Intracellular NADH exists in both free and protein-bound forms, which can be distinguished by their distinct fluorescence lifetimes (∼400 ps and ∼2500 ps, respectively)^20,21^. Changes in glycolytic flux, TCA cycle activity, or electron transport chain function alter the relative contribution of these pools. Oxidation of free NADH to non-fluorescent NAD^+^ reduces the relative contribution of the short-lifetime NADH pool and increases the proportional contribution of protein-bound NADH. Given the distinct fluorescence lifetimes of free and protein-bound NADH, this shift is reflected in an increase in the mean fluorescence lifetime. Such changes in mean lifetime have been widely used as a proxy for alterations in the NAD^+^/NADH balance and cellular metabolic state. They can be monitored non-invasively using fluorescence lifetime imaging microscopy (FLIM), as previously demonstrated in immune cells, stem cells, and cancer cells^20–26^.

It should be noted that the autofluorescence signal detected in NADH FLIM measurements includes contributions from both NADH and NADPH, whose fluorescence lifetimes cannot be readily distinguished^27^. In particular, free NADH and free NADPH exhibit similar fluorescence lifetimes, and an oxidation of free NADPH can likewise result in a shift toward longer mean lifetimes. Accordingly, while we refer to changes in the NAD^+^/NADH balance throughout this work, variations in the NADP^+^/NADPH pool may also contribute to the observed lifetime changes and should be considered in the interpretation of the data.

TGF-β1 has been shown to modulate NADH dynamics through activation of NAD(P)H oxidases, leading to increased reactive oxygen species production and subsequent induction of α-SMA^28^. These signaling-driven metabolic shifts suggest that NADH fluorescence lifetimes can report CAF activation state. Building on this coupling between signaling, metabolism, and phenotype, we show that NADH FLIM enables a reliable and quantitative, label-free, non-destructive read-out of CAF activation, allowing real-time monitoring of metabolic remodeling and complementing conventional marker-based approaches.

## Material and methods

### Cell culture

The KM12L4a cell line was kindly provided by the laboratory of Dr. I. Fidler (MD Anderson Cancer Center) and modified to express mScarlet for selection and visualization in co-culture experiments. Stable KM12L4a-mScarlet cells were established by lentiviral transduction of KM12L4a cells with Lyn-mScarlet lentiviruses. Lentiviruses were made by co-transfecting HEK-293T cells with a pHR vector containing the Lyn-mScarlet sequence and with envelope plasmid pMD2 and packaging plasmid pCMV. pCMVR8.74 (Addgene plasmid #22036) and pMD2.G (Addgene plasmid #12259) were a gift from Didier Trono. The three plasmids were mixed (ratio pHR:packaging:envelope, 1:2:1.2) and Turbofect (Thermofisher) was used to transfect the cells. Two days post-transfection, viral particles were collected, filtered to removed cell debris and added onto the KM12L4a cells. KM12L4a transfected cells were selected based on their resistance to blasticidin and further expanded. Isolation, characterization and culture of human colorectal CAF were previously described^29,30^.

All cells were cultured in Dulbecco’s Modified Eagle Medium (Gibco™ DMEM, ThermoFisher Scientific), supplemented with 5% Gibco GlutaMAX™, 10% fetal bovine serum (Gibco™ FBS), and gentamicin (1:1000). To select for cells expressing mScarlet, which was cloned with a Blasticidin resistance marker, 10 µg/mL Blasticidin S was added to the medium. Cells were cultured following standard procedures at 37°C with 5% CO_2_ and passaged at confluency using TrypLE™ Express (ThermoFisher Scientific, cat. no. A1217702).

### CAF activation

To ensure robust activation of cancer-associated fibroblasts (CAFs), cells were seeded at high density on collagen-coated surfaces and treated with TGF-β. A 24-well glass bottom plate was coated with 400µL Collagen type I (Sigma Aldrich, Cat. No. CLS354236) in acetic acid (ACROS organics, Cat. No. 148930010) (7.5µg/mL). The plate was incubated at 37°C for at least 20 minutes to allow coating, followed by three washes with 500 µL of phosphate-buffered saline (PBS) (Gibco™ PBS 10X, Cat. No. 70-011-044, ThermoFisher Scientific).

CAFs were then split, counted, and seeded at a density of 50 000 cells per well onto the coated wells and incubated overnight at 37 °C. The next day, the culture medium was replaced with starvation medium (DMEM without supplements), and the cells were incubated for an additional 24 hours. Fibroblast activation was induced by replacing the starvation medium with supplemented DMEM containing 2 ng/mL TGF-β (in 4mM HCl (Honeywell) and 0.1% BSA) (Gibco™, BSA fraction V 7.5%, Cat. No. 15260037, ThermoFisher Scientific) and incubated for 48h.

Control, non-activated CAFs were seeded at a lower density (10,000 cells per well) on uncoated wells and incubated for 24 h prior to imaging.

### Immunostaining

To assess activation of CAFs, cells were fixed in 4% paraformaldehyde (ThermoFisher Scientific, v/v 16% in PBS) for 15 minutes at room temperature. Subsequently, cells were permeabilized using 0.1% Triton X-100 and 0.1% Tween-20 (Merck, Cat. No. P7949-500ML) in PBS for 15 minutes and blocked for 1 hour using a blocking solution (10% fetal bovine serum (FBS) and 0.1% Tween-20 in PBS), to prevent nonspecific antibody binding. Cells were incubated overnight at 4 °C with a primary goat anti-α-SMA antibody (Merck, Cat. No. A5228-200UL) (1:400), followed by incubation with an Alexa Fluor 647– conjugated secondary antibody (1:1000). This assay served to confirm whether cells displayed elevated α-SMA expression consistent with activation.

### Western blot

Cells were lysed on ice in RIPA buffer (Carl Roth) with protease inhibitors (1:100) (MedChemExpress, Cat. No. HY-K0010). Lysates were centrifuged for 15 min at maximum speed at 4 °C, and protein concentrations were determined using the BCA method^31^. Equal amounts of protein extract (10 µg) were mixed with Laemmli sample buffer and denatured prior to electrophoresis.

Proteins were separated on 10% SDS–PAGE gels at 150V for 45 minutes and transferred to nitrocellulose membranes at 25V for 90 minutes. Membranes were blocked for 1 h in PBS-T (0.1% Tween-20) and 3% BSA (Sigma Aldrich, Cat. No. A3294-10G), followed by incubation overnight at 4 °C with primary anti-α-SMA antibody (1:1000) (Merck, Cat. No. A5228-200UL) and anti-tubulin antibody (1:1000) (Proteintech, Cat. No. 11224-1-AP)). After washing with PBS-T, membranes were incubated for 2h at room temperature with ALEXA488- (Proteintech Cat. No. SA00013-2) and ALEXA647-conjugated secondary (ThermoFisher Scientific, Cat. No. A21235) antibodies and was visualized using an iBRIGHT imager (ThermoFisher Scientific).

### NADH measurements

Fluorescence lifetime imaging microscopy (FLIM) was performed to assess activation-associated metabolic changes by measuring shifts in the mean NADH fluorescence lifetime, reflecting changes in the free-to-bound NADH ratio. Fluorescence decay curves were analyzed in LAS X software using a biexponential decay model corresponding to free NADH (∼400 ps) and protein-bound NADH (∼2500 ps). The goodness of fit was evaluated using the χ^2^ value, with values between 0.7 and 2 considered indicative of an acceptable fit^32^. Accurate lifetime fitting required sufficient photon counts (>700,000), as low signal levels can adversely affect fitting reliability. Mean fluorescence lifetime was calculated using an intensity-weighted approach. Cellular compartments (mitochondria, nucleus, and cytosol) were identified based on NADH autofluorescence intensity. Mitochondria were segmented based on intensity, nuclear regions were manually selected within clearly identifiable cell nuclei, and the cytosol was defined as the remaining cellular area excluding mitochondria and nuclei.

During live-cell NADH-FLIM acquisition, individual imaging positions were recorded to allow direct correlation with downstream analyses. Following FLIM measurements, the sample was fixed and subjected to immunofluorescence staining for α-SMA as described above, enabling validation of fibroblast activation status in the same cells that were previously analyzed by FLIM.

### Wound healing assay

To assess cell migration in a dynamic setting, KM12L4A cancer cells expressing mScarlet and CAF cells were seeded in separate compartments of a two-well Ibidi silicone insert (Cat. No. 80209) placed in 24-well plates. For each condition, 70 µL of cell suspension was added to each compartment: 70 000 non-activated CAFs on one side and 150 000 KM12L4A cancer cells on the other.

Cells were allowed to attach overnight, after which the inserts were removed to generate a defined cell-free gap between the two cell populations. Subsequently, starvation medium was added to each well to limit proliferation and allow analysis of cell migration and CAF activation. CAF activation was monitored by FLIM at approximately 24h intervals, focusing on CAFs located close to the cell-free gap.

### Fluorescence microscopy

Multiphoton FLIM and confocal fluorescence imaging were performed on a Leica TCS SP8 DMi8 inverted microscope (Leica Microsystems, Germany). The system is equipped with a tunable femtosecond multiphoton laser (Mai Tai DeepSee, Spectra Physics; 690–1040 nm), a white light laser (WLL, 470–670 nm), and hybrid detectors (HyD) for high-sensitivity detection. Imaging was carried out using a 63× water-immersion objective (HC PL APO, NA 1.2).

For FLIM experiments, NADH autofluorescence was excited at 730 nm using the multiphoton laser with 5.75 mW laser power at the objective. Fluorescence emission was collected through a 440/80 nm band-pass filter (F47-441) in combination with a 494 nm long-pass beam splitter (F48-908) (AHF Analysentechnik). Lifetime data were acquired using the Leica FALCON FLIM module, with a detector gain of 60, a pixel dwell time of 15.3875 µs, and three frame accumulations per image to ensure sufficient photon counts while minimizing phototoxicity.

During FLIM acquisition, imaging positions were recorded using the Navigate (Mark and Find) function in LAS X software, allowing precise relocation of the same cells at later time points. Following FLIM measurements, samples were fixed and subjected to immunostaining. The previously recorded positions were then revisited, and the same cells were imaged using confocal laser microscopy.

For immunofluorescence imaging, excitation was performed using the white light laser tuned to 647 nm with 2.05 µW laser power at the objective. Fluorescence emission was detected in the range of 644– 707 nm using standard HyD detectors with an acousto-optical beam splitter (AOBS) and freely selectable emission windows. The pixel dwell time was 7.6875 µs.

The microscope was fully enclosed and equipped with an environmental control system (The Cube chamber heater and The Brick CO_2_ gas mixer (Life imaging Services)) to minimize disruption of cell dynamics. All FLIM measurements were acquired and processed using the FLIM module of the LAS X software (Leica Microsystems)

## Results & discussion

### Validation of TGFβ-induced CAF activation

To establish a controlled model of CAF activation for downstream metabolic analysis, fibroblasts were treated with the well-established activator TGF-β, which induces a classical activated CAF phenotype^7^. α-SMA, a widely accepted hallmark of the myofibroblast-like, contractile CAF phenotype,^9,10^ was used as a reference marker to validate CAF activation under our experimental conditions.

CAF activation was first confirmed at the population level by Western blot analysis of α-SMA expression (Fig. 1A). Following TGF-β treatment, CAFs displayed a clear increase in α-SMA protein abundance compared to untreated controls, confirming successful induction of an activated state.

**Figure 1.**
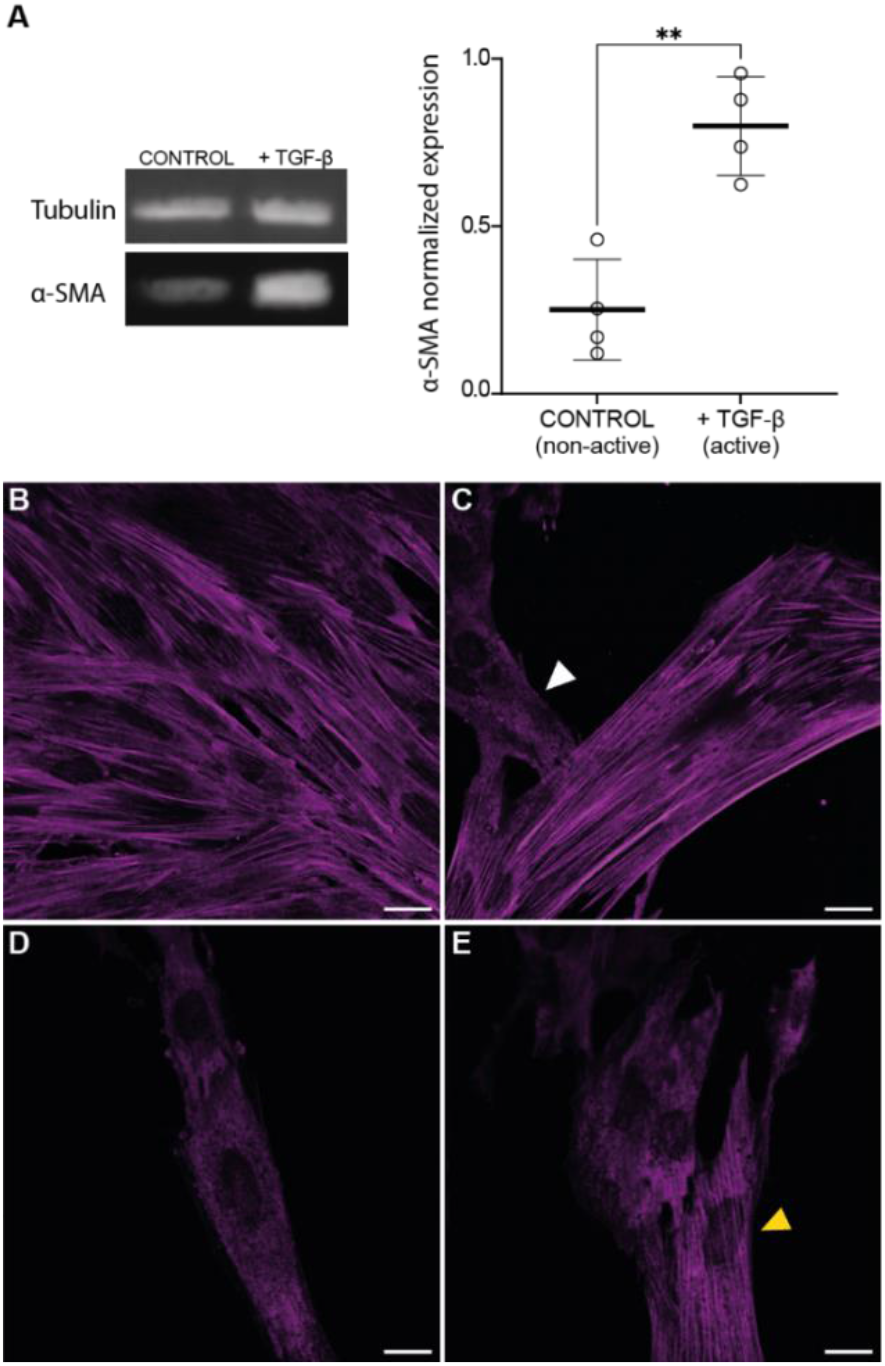
**A**. Western blot confirming increased α-SMA expression in TGF-β–treated CAFs compared to untreated controls. B-C. α-SMA immunostaining of TGF-β–treated CAF cells showing predominantly activated cells with robust, fiber-like α-SMA organization (B), alongside occasional cells lacking clear α-SMA stress fiber formation (C, white arrow), illustrating residual heterogeneity within the activated population. D-E. non-activated cells without detectable α-SMA fiber organization (D), with rare cells displaying partial α-SMA fiber-like structures (E, yellow arrow). Scalebars: 20µm.

Because CAF populations are heterogeneous, increased α-SMA levels detected by Western blot do not necessarily reflect uniform activation across all cells. α-SMA expression was therefore additionally examined at the single-cell level using immunofluorescent staining (Fig. 1B-E). TGF-β–treated CAFs predominantly exhibited robust and well-organized αSMA stress fibers, with the majority of cells (∼90%) displaying this myofibroblast-like, contractile phenotype. In contrast, untreated CAFs largely showed a non-activated morphology without detectable αSMA fiber formation, although occasional cells exhibited some αSMA organization, potentially reflecting local cell-cell interactions or activation through close proximity.

Together, these imaging and biochemical validations confirm successful CAF activation and provide a robust biological framework for interpreting the metabolic changes assessed by NADH FLIM in the subsequent experiments.

### NADH FLIM as a marker of CAF activation

To determine whether TGF-β–induced CAF activation is accompanied by detectable changes in cellular NAD^+^/NADH balance, we next evaluated if NADH fluorescence lifetimes differ between activated and non-activated CAFs using FLIM. Of note, given the heterogenous character of CAF activation, only cells confirmed to be activated based on α-SMA expression were included in the analysis (Fig. 2A).

**Figure 2.**
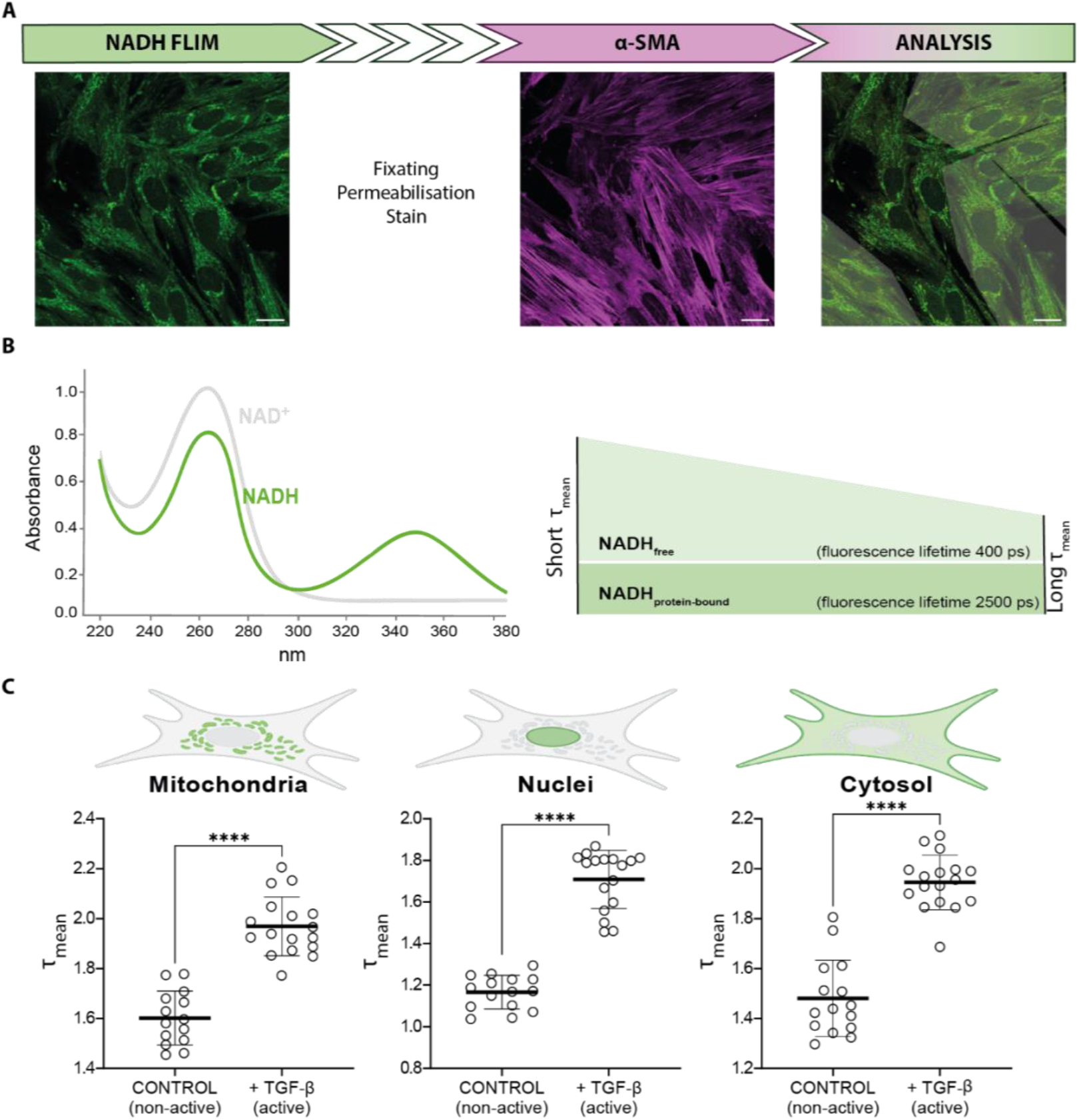
**A**. Experimental workflow illustrating the analysis strategy. Live cells were first imaged by NADH fluorescence lifetime imaging microscopy (FLIM) to assess intrinsic autofluorescence, followed by fixation, permeabilization and α-smooth muscle actin (α-SMA) immunostaining to identify activated CAFs for analysis (selected shadow regions). Scale bars: 20µm B. Schematic overview of NADH fluorescence lifetime principles, illustrating the spectral properties of NAD^+^/NADH, the distinction between free (short lifetime) and protein-bound (long lifetime) NADH pools, as well as how a decrease in the amount of free NADH results in an increase in the mean fluorescence lifetimes (τ_mean_). C. Quantification of mean NADH lifetimes (τ_mean_) in mitochondria, nuclei and cytosol, showing increased lifetimes in activated CAFs compared to non-activated controls. Each data point represents the mean lifetime recorded for all active or nonactive cells in one field of view, with four replicates.

As mentioned in the introduction, the NADH autofluorescence signal originates from reduced NADH, whereas oxidized NAD^+^ is non-fluorescent in the relevant spectral range (Fig. 2B). Cellular NADH exists in two pools: free, dynamic NADH with a short fluorescence lifetime (∼400 ps) and more stable protein-bound NADH with a longer lifetime (∼2500 ps)^20,21^. We hypothesized that the metabolic reprogramming associated with TGF-β–induced CAF activation will promote the consumption and oxidation of free NADH, thereby shifting the balance toward protein-bound NADH and resulting in an increase in the mean fluorescence lifetime (τ_mean_, Fig. 2B).

As shown in Figure 2A, NADH autofluorescence is detected throughout the cell, with the highest signal intensity typically observed in mitochondria, reflecting the higher concentration of NADH in this compartment. To evaluate whether TGF-β activation induces compartment-specific metabolic changes or a more global shift in NADH dynamics, mean fluorescence lifetimes (τ_mean_) were quantified separately in mitochondrial, nuclear, and cytosolic regions (Fig. 2C).

As expected, baseline τ_mean_ values differed between cellular compartments, reflecting differences in local NADH dynamics. Comparable lifetimes were observed in mitochondria and cytosol (1.60 ± 0.11 ns and 1.48 ± 0.15 ns, respectively), whereas lower τ_mean_ values were detected in the nucleus (1.17 ± 0.08 ns, Fig 2C). Despite these compartment-specific baseline differences, activated CAFs exhibited a consistent shift toward longer mean NADH lifetimes across all three compartments compared to non-activated controls (Fig. 2C), with τ_mean_ values of 1.97 ± 0.12 ns in mitochondria, 1.71 ± 0.14 ns in the nucleus, and 1.95 ± 0.11 ns in the cytosol. Importantly, the direction and magnitude of this lifetime shift were comparable between compartments, indicating that TGF-β–induced changes in NADH dynamics reflect a global alteration in cellular metabolic state rather than a compartment-restricted effect.

These results support a model in which TGF-β activation induces metabolic reprogramming in CAFs, characterized by increased glycolytic and catabolic activity and enhanced consumption of free NADH during the production of energy-rich metabolites such as L-lactate and pyruvate^12–14,28^. Oxidation of free NADH to NAD^+^ reduces the free NADH pool and increases the relative contribution of the longer-lived, protein-bound NADH fraction, thereby shifting τ_mean_ toward longer lifetimes.

Our results demonstrate that the shift toward longer NADH fluorescence lifetimes provides a robust metabolic signature of CAF activation (Fig. 2C). However, it is important to note that previous research has shown that some cultured cells can undergo substantial metabolic changes following cryopreservation and thawing, such that cells thawed from different vials may display distinct metabolic states, even when originating from the same donor batch^33–35^. Because the intracellular free-to-bound NADH ratio reflects the underlying metabolic state, such thawing-induced variability can lead to differences in absolute NADH fluorescence lifetime values between cell populations. In addition, CAFs are inherently heterogeneous and comprise multiple distinct subpopulations with distinct metabolic profiles^4,12,14,36^. Moreover, activated CAF comprise different phenotypes, namely myCAF and iCAF phenotypes, further contributing to the population-dependent variability in absolute NADH lifetimes^6^.

Consistent with these considerations, CAFs grown from an additional vial originating from the same donor batch but thawed and cultured at a different time exhibit different baseline τ_mean_ values. Nevertheless, despite these differences in absolute lifetimes, TGF-β activation induced a comparable shift in τ_mean_ between activated and corresponding control conditions, with a similar direction and magnitude of change (Supplementary Figure S1). Accordingly, the use of NADH FLIM as a readout of CAF activation requires paired, parallel measurements of activated and control samples acquired under identical experimental conditions.

Together, these results highlight the utility of NADH FLIM as a sensitive, dynamic, and label-free readout of CAF activation, while emphasizing the importance of biological controls for accurate interpretation.

### Dynamic Monitoring of CAF Activation

To assess whether NADH-FLIM can dynamically report CAF activation in a physiologically relevant context, we performed a wound-healing assay that mimics tumor-stroma interactions during cell migration. In this assay, CAFs were seeded on one side of a well-defined cell-free gap and KM12L4A colorectal cancer cells were placed on the opposite side. The presence of cancer cells is essential in this assay, as tumor-derived biochemical cues are known to activate CAFs and stimulate their migration toward the tumor compartment. This configuration therefore allows testing, in a dynamic setting, whether NADH lifetime changes reflect this activation process in real time.

During the assay, CAFs located at or migrating into the wound edge were selected for analysis, as these cells are most directly exposed to tumor-derived signals and actively engaged in the activation and migration process. Representative NADH fluorescence images acquired during the assay are shown in Fig. 3A, with cells selected for lifetime analysis indicated by shaded regions. Because NADH is most abundant in mitochondria, and comparable activation-induced τ_mean_ shifts were observed across all cellular compartments, mitochondrial regions were selected for further analysis. Focusing on the compartment with the highest fluorescence intensity enabled the collection of sufficient photon counts over shorter acquisition times, thereby reducing phototoxicity during live-cell imaging.

**Figure 3.**
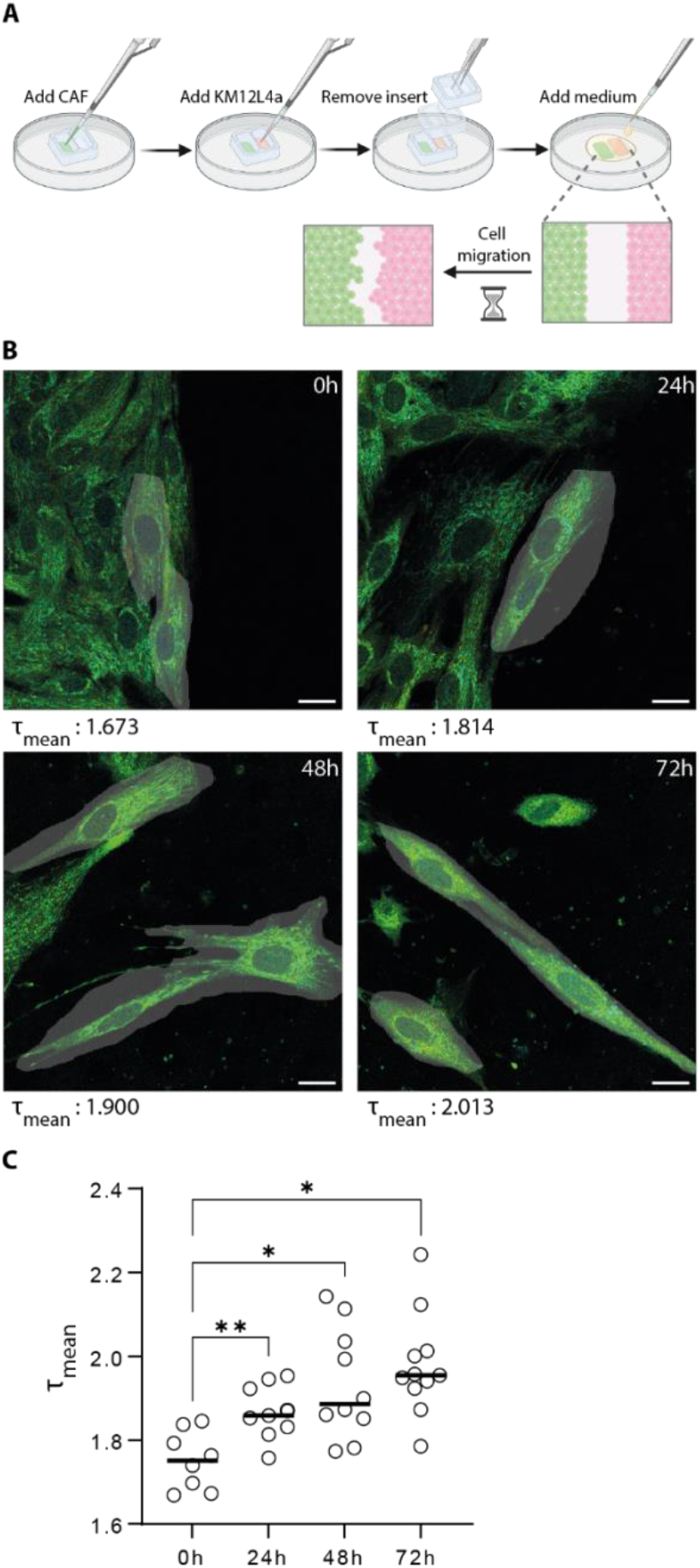
**A**. Schematic overview of the experimental wound healing assay. NADH-FLIM imaging is subsequently performed at defined time points to monitor metabolic changes during cell migration. B. NADH-FLIM images (green: NADH autofluorescence) of CAFs located at or migrating into the cell free gap at 0, 24, 48, and 72 hours. Cells selected for mitochondrial lifetime analysis are shown in grey, and their corresponding mitochondrial mean lifetimes are indicated for each time point. Scalebars: 20µm C. Quantification of mitochondrial NADH mean lifetimes over time. A significant increase is observed already at 24h compared to 0h, with lifetimes continuing to rise at 48h and peaking at 72h. Each data point represents cells selected within the field of view close to the gap, as illustrated in panel B; three replicates were analyzed.

Quantification of mitochondrial mean NADH lifetimes revealed a progressive increase over time (Fig. 3B). Already at 24h, τ_mean_ values were significantly elevated compared to baseline (0h), increasing from 1.75 ± 0.07 ns at 0 h to 1.87 ± 0.06 ns at 24 h, indicating an early CAF activation. Mean lifetimes continued to rise at 48h and peaked at 72h, with average τ_mean_ values of 1.98 ± 0.12 ns. Notably, the variability in τ_mean_ increased at later time points, reflected by a broader distribution of lifetimes, suggesting increasing metabolic heterogeneity among migrating CAFs as activation progressed (Fig. 3B).

Together, these data demonstrate that NADH-FLIM sensitively captures the temporal evolution of metabolic remodeling associated with CAF activation during migration in a live-cell context.

## Conclusions

In this study, we demonstrate that NADH fluorescence lifetime imaging microscopy provides a robust, label-free readout of CAF activation by reporting activation-associated metabolic remodeling. TGF-β– induced activation resulted in a consistent shift toward longer NADH lifetimes across cellular compartments, while dynamic wound-healing assays revealed that NADH-FLIM can track the temporal progression of CAF activation in live, migrating cells. Importantly, while absolute lifetime values vary due to biological and technical heterogeneity, the relative lifetime shift associated with activation is preserved when analyzed in paired, parallel conditions. Together, these findings establish NADH-FLIM as a powerful approach for real-time, non-invasive monitoring of CAF activation dynamics in physiologically relevant settings.

## Acknowledgements

We acknowledge additional financial support from Research Foundation of Flanders (FWO) research grant (G0C2422N), postdoctoral fellowship for GS (12AML24N), Foundation against Cancer (2024-151) and from the KU Leuven (IDN/20/021 and C14/22/085).

## Notes

### Competing Interest Statement

The authors have declared no competing interest.

