## Supporting Figure 1 for "Label-Free Monitoring of Cancer-associated Fibroblast Activation using NADH Fluorescence Lifetime Imaging"

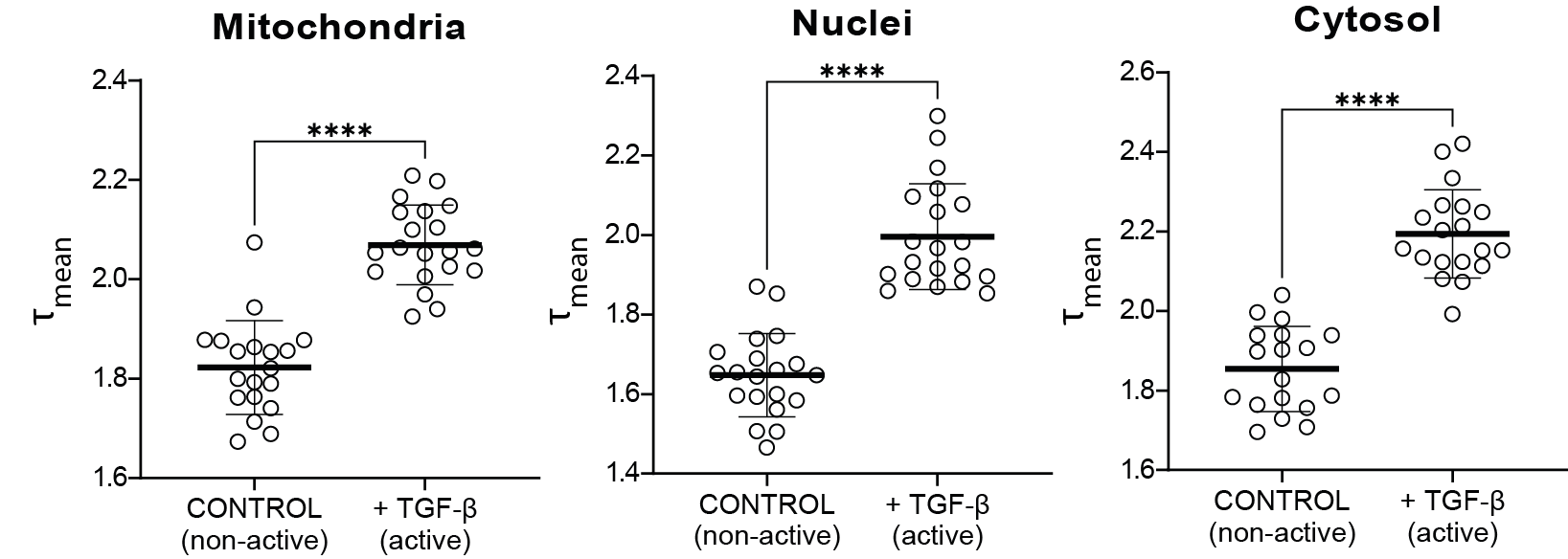

**Supplementary Figure S1.** Quantification of mean NADH fluorescence lifetimes (τ_mean_) in mitochondria, nuclei, and cytosol of control and TGF-β–activated CAFs. Data are shown for CAFs cultured from an additional vial originating from the same donor batch but thawed and expanded at a different time. Upon TGF-β activation, τ_mean_ increased across all compartments, from 1.82 ± 0.09 ns to 2.07 ± 0.08 ns in mitochondria, from 1.65 ± 0.10 ns to 2.00 ± 0.13 ns in nuclei, and from 1.85 ± 0.11 ns to 2.19 ± 0.11 ns in the cytosol. Each data point represents the mean lifetime recorded for all active or nonactive cells in one field of view.
